# Optical metabolic imaging identifies metabolic shifts and mitochondria heterogeneity in POLG mutator macrophages

**DOI:** 10.1101/2025.01.13.632822

**Authors:** Linghao Hu, A. Phillip West, Alex J. Walsh

**Affiliations:** Department of Biomedical Engineering, Texas A&M University, 3120 TAMU, College Station, TX, 77843, USA; The Jackson Laboratory, Bar Harbor, ME, 04609, USA

## Abstract

The polymerase gamma (POLG) gene mutation is associated with mitochondria and metabolism disorders, resulting in heterogeneous responses to immunological activation and posing challenges for mitochondrial disease therapy. Optical metabolic imaging captures the autofluorescent signal of two coenzymes, NADH and FAD, and offers a label-free approach to detect cellular metabolic phenotypes, track mitochondria morphology, and quantify metabolic heterogeneity. In this study, fluorescence lifetime imaging (FLIM) of NAD(P)H and FAD revealed that POLG mutator macrophages exhibit a decreased NAD(P)H lifetime, and optical redox ratio compared to the wild-type macrophages, indicating an increased dependence on glycolysis. FLIM revealed that both wild-type and POLG mutator macrophages switch to a decreased NAD(P)H *τ*_*1*_, and *τ*_*m*_ after immune stimulation by Lipopolysaccharides (LPS). Furthermore, a bimodality index of subpopulation analysis identified heterogenous populations of POLG mutator macrophage responses under immune challenge by LPS. Moreover, to quantify the mitochondria variations in POLG mutator macrophages, a customized thresholding image processing pipeline was developed to segment mitochondria regions within each cell from the NADH image, allowing for the feature analysis of mitochondria clusters. Consequently, the wild-type macrophages exhibited a higher percentage of mitochondria-containing pixels and longer lengths of connected mitochondria, as compared with POLG mutated macrophages. Altogether, these results illustrate the potential of optical metabolic imaging for non-invasive detection and quantification of cellular metabolism, metabolic heterogeneity within cell populations, and intra-cellular mitochondria morphology differences in POLG mutator macrophages. Optical metabolic imaging will be valuable for studying POLG-mutation diseases and evaluating efficacy of potential therapies.

## Introduction

The polymerase gamma (POLG) gene encodes polymerase gamma (pol γ), a DNA polymerase crucial for DNA replication and repair [1, 2]. Specially, pol γ is the only known DNA polymerase that is active within mitochondria to facilitate mitochondria DNA (mtDNA) replication and repair [1]. Therefore, the mutations of POLG serve as a locus for several mitochondrial diseases, such as primary mitochondrial diseases (MD), parkinsonism, cancer, progressive external ophthalmoplegia (PEO), sensory ataxic neuropathy dysarthria and ophthalmoparesis (SANDO), and Alpers-Huttenlocher syndrome (AHS) [1, 3, 4]. The POLG mutation destroys mitochondria function and impairs OXPHOS metabolism leading to an increased reliance on glycolysis. Furthermore, POLG mutations can lead to heterogeneity in cell populations due to mitochondrial heteroplasmy [5], resulting in diverse effects and various immune responses among cell populations. Aberrant activation of the type I interferon (IFN-I) signaling has been observed in POLG mutator cells, leading to enhanced proinflammatory cytokine responses, and accelerated metabolic dysfunction [6]. Thus, non-invasive single-cell techniques to detect metabolic changes under immunological activation and identify mitochondria in live cells could significantly advance our understanding of how POLG mutations shape immune responses and mitochondria dysfunctions.

Optical metabolic imaging (OMI) detects metabolic information of cells in a label-free modality with subcellular spatial resolution. When integrated with cell segmentation techniques, OMI resolves single-cell information and heterogeneity within populations of cells [7]. OMI measures the autofluorescence intensity and lifetimes of two metabolic co-enzymes: reduced nicotinamide adenine dinucleotide (NADH) and flavin adenine dinucleotide (FAD) [8, 9]. Both NADH and FAD facilitate oxidation-reduction reactions in various metabolic pathways. Specifically, NADH is an electron acceptor in glycolysis and an electron donor in oxidative phosphorylation, whereas FAD is an electron donor in oxidative phosphorylation. The oxidized form of NADH, NAD^+^, does not exhibit fluorescence, and similarly, the reduced form of FAD, FADH_2_, is non-fluorescent. Therefore, the ratio of NADH and FAD intensity values, a metric termed the optical redox ratio, is sensitive to the redox state of cells and tissues [9-11]. NADH and its phosphorylated form, NADPH, have the same fluorescence spectral properties, so NAD(P)H is used to represent the measured signal from cells which includes contributions from both NADH and NADPH [12].

The fluorescence lifetime measures the duration a fluorophore remains at the excited state before returning to the ground state, and the fluorescence lifetimes of NADH and FAD are sensitive to the protein binding activities of the coenzymes, providing additional information related to metabolic activity. Protein-bound NADH exhibits a longer lifetime than the free NADH, due to self-quenching of the free molecule [13]. Conversely, free FAD has a longer lifetime than the protein-bound FAD [14]. In summary, the fluorescence intensity and lifetime of NADH and FAD offer a functional measure of metabolic activity in individual cells or whole tissue sections and have been used to identify metabolic differences in precancerous tissues [15], detect T-cell activation [16], predict the efficiency of stem cell differentiation [17], and study drug treatment response of cancer cells [18].

As NADH exists at high concentrations within mitochondria [19], spatial variations in NADH fluorescence intensity within cells can reveal intracellular mitochondrial organization. Quantification of the power spectral density of NAD(P)H intensity images reveals mitochondria subcellular patterns and has been applied to resolve differences in spatial NAD(P)H patterns between normal and diseased tissue [20, 21]. Recent studies have leveraged the spatial patterns within NAD(P)H images by using convolutional neural networks to discriminate metabolic phenotypes in cancer cells and T cells [22, 23]. Furthermore, high-resolution autofluorescence imaging of NAD(P)H reveals brighter pixels in mitochondria regions compared to cytosol regions and enables the visualization of mitochondria networks using high magnification and numerical aperture objectives [24, 25]. Furthermore, several novel software tools have been developed for the analysis of mitochondrial networks including mitometer [26], MINA [27], and Mitochondria Analyzer [28]. These tools are designed to segment and track spatiotemporal mitochondria in live-cell fluorescent images, enabling the identification of alterations in mitochondrial morphology, motion, and fission and fusion dynamics. While these software methods have been developed and demonstrated for mitochondria labeled with fluorescent probes, their applicability to NAD(P)H images of macrophages is limited due to the low contrast of autofluorescence confined nature within small cells.

In this study, autofluorescence lifetime imaging of NAD(P)H and FAD was used to detect the metabolic responses of wild-type (WT) and POLG-mutation primary bone marrow-derived macrophages (BMDMs) upon lipopolysaccharide (LPS) stimulation. Furthermore, the bimodality index (BI) of three key optical metabolic parameters, NAD(P)H *τ*_*m*_, FAD *τ*_*m*_, and the optical redox ratio revealed heterogeneity within the unstimulated and LPS-challenged POLG population. A convergence in metabolism between WT and POLG BMDMs over successive generations was observed through the projection of OMI features using unsupervised machine learning algorithms. In autofluorescence images of BMDMs, mitochondria tend to cluster together, making it challenging to resolve their network structure even under high magnification (100X). This limits the application of existing software to perform mitochondria segmentation, morphology, and network assessment. To address this, we developed a customized image-processing pipeline that identifies mitochondria regions within cells, segments mitochondria clusters, and quantifies the skeleton length of these regions. As a result, WT BMDMs occupy a larger percentage of the mitochondria area and exhibit longer mitochondria skeletons than the POLG BMDMs. In summary, this label-free technique enables the detection of metabolism and mitochondria in POLG mutator cellular models, enhancing our understanding of metabolic heterogeneity, variations, and potential chemotherapy responses.

## Methods

### POLG and WT BMDM extraction, culture, and LPS treatment

The WT and POLG immortalized BMDMs were created by Y.J. Lei, and A.P. West, as fully described in [6, 29]. C57BL/6J and POLGD257A/D257A mutator mice were purchased from the Jackson Laboratory and used to generate POLG +/+ and POLG D257A/D257A experimental mice. Primary BMDMs were isolated from femur and tibia bones, and immortalized cultures were created by infecting the primary macrophages with J2 recombinant retrovirus. The WT and POLG BMDMs were cultured in DMEM (gibco, 11965-092) supplemented with 10% fetal bovine serum (FBS, CORNING, 35-011-CV). Both the younger generation and the older generation of POLG and WT BMDMs originated from the same cell source. The younger BMDMs underwent approximately 7-10 passages and were then frozen and stored in a -80 ^°^C freezer for approximately 3 months. Subsequently, the older BMDMs were obtained by thawing the previously frozen cells, culturing them, and finally reaching passages 35 to 40. A density of 10^5 cells BMDMs were seeded per 35mm glass-bottom imaging dish 48 hours before imaging in normal growth media. Lipopolysaccharide (LPS) (5 μg/ml, eBioscience, 00-4976-93) was added 24 hours before imaging for the LPS-stimulated groups. To ensure the reliability of the results, three experimental replicates were performed, with a total of three dishes prepared for each group. To evaluate the correlation between NADH intensity and mitochondria localization, in a subset of imaging dishes, MitoTraker Deep Red (Cell Signaling, #8778) was added to the media (50 nM) and incubated for 30 minutes at 37ºC to label the mitochondria regions.

### Fluorescence lifetime and intensity imaging

NAD(P)H and FAD autofluorescence intensity and lifetime images of BMDMs were captured using a customized multiphoton imaging system (Mariana, 3i) coupled with a time-correlated single-photon counting (TCSPC) electronics module (SPC -150N, Becker & Hickl). The fluorescence of NAD(P)H and FAD was excited using a tunable (680 -1080nm) Ti: sapphire femtosecond laser (COHERENT, Chameleon Ultra II) and detected using photomultiplier tubes (PMT, HAMAMATSU, H7422PA-40) with bandpass filters: 447/60 nm for NAD(P)H, and 550/88 nm for FAD. Specifically, NAD(P)H was excited at 750 nm with a power of 7.2 mW, and FAD was excited at 890 nm with a power of 5.5 mW. A 100x objective (1.46 NA) was used for fluorescence lifetime imaging of NAD(P)H and FAD, and each fluorescence lifetime image (256 × 256 pixels, 108 × 108 μm^2^) was acquired with a pixel dwell time of 50 μs and 3 frame repeats. For fluorescence intensity imaging, we used an additional 150x objective (1.35 NA) to capture NAD(P)H fluorescence images. Each intensity image with a dimension of 1024 × 1024 pixels (108 × 108 μm^2^ for 100X and 72 × 72 μm^2^ for 150x) was acquired with a pixel dwell time of 2.0 μs and 1 frame repeat. The instrument response function (IRF) was measured by the second harmonic generated signal of urea crystals, which were excited at 900 nm and detected using the NAD(P)H channel. MitoTracker fluorescence was excited at 800 nm and detected using a PMT (HAMAMATSU, C6270) with a bandpass filter of 675/67 nm. Both NAD(P)H and FAD fluorescence images were captured in at least five randomly selected positions for each dish. The NAD(P)H and FAD fluorescence lifetime images of the younger generation of BMDMs were collected by L.H. Hu, and A.J. Walsh using the same microscope with 40x objective (1.1 NA), and the imaging details are fully covered in Hu et al. [29].

### Fluorescence lifetime analysis of each cell

The fluorescence lifetime decays were analyzed by SPCImage (Becker & Hickl). For both NAD(P)H and FAD fluorescence lifetime images, a binning of nine surrounding pixels was used, and the decay curve of each pixel was deconvoluted from the measured IRF and then fitted to a bi-exponential model, (*It*) =α_1_*e*^−*t/*τ_1_^ + α_2_*e*^−*t/*τ_2_^ + *C*, where *I(t)* represents the fluorescence intensity as a function of time *t, τ*_*1*_, and *τ*_*2*_ are the short and long lifetimes, *α*_*1*_ and *α*_*2*_ are their corresponding fractions and *C* accounts for an offset due to background light. Single-cell segmentation was based on the NAD(P)H intensity image and achieved using Cellpose [30, 31]. Different cell diameters were set depending on the objective used to acquire the image, diameter = ∼100 pixels for images taken with the 100x objective, and ∼130 pixels for images taken with the 150x objective. The CP model in Cellpose [31] was used to obtain the segmented mask of each cell. The number of cells in each group segmented with Cellpose is summarized in Sup. Table 1.

Image processing was performed in MATLAB to calculate the redox ratio image (FAD/ (FAD + NAD(P)H)), and the weighted average NAD(P)H and FAD fluorescence lifetime images (*τ*_*m*_ *= α*_*1*_*τ*_*1*_ *+ α*_*2*_*τ*_*2*_). The cytoplasmic region of each cell was separated from the cellular mask by using an intensity threshold of 40% of the maximum NAD(P)H intensity of that cell. Autofluorescence lifetime endpoints for each cell were then obtained by computing the mean and standard deviation of pixel values within the segmented cytoplasmic regions across lifetime images. The fluorescence lifetime data of the younger BMDMs were obtained from Hu et al. [29].

### Mitochondria segmentation and feature calculation

Segmentation of mitochondria was achieved using a customized MATLAB image processing pipeline. Each step of the segmentation pipeline is illustrated for an example cell image in Figure 1. For the segmented NAD(P)H intensity image of each cell, a 2D Gaussian filter with a standard deviation of 1 was applied to smooth the image and filter random noise. Then, an empirical threshold, set at half of the maximum NAD(P)H intensity pixel values of that cell, was used to extract the brighter pixel regions. As mitochondria appear as clusters of bright pixels in the NAD(P)H intensity images of BMDMs, a threshold of 50% of the maximum intensity within a cell was used to extract the mitochondria regions.

**Figure 1.**
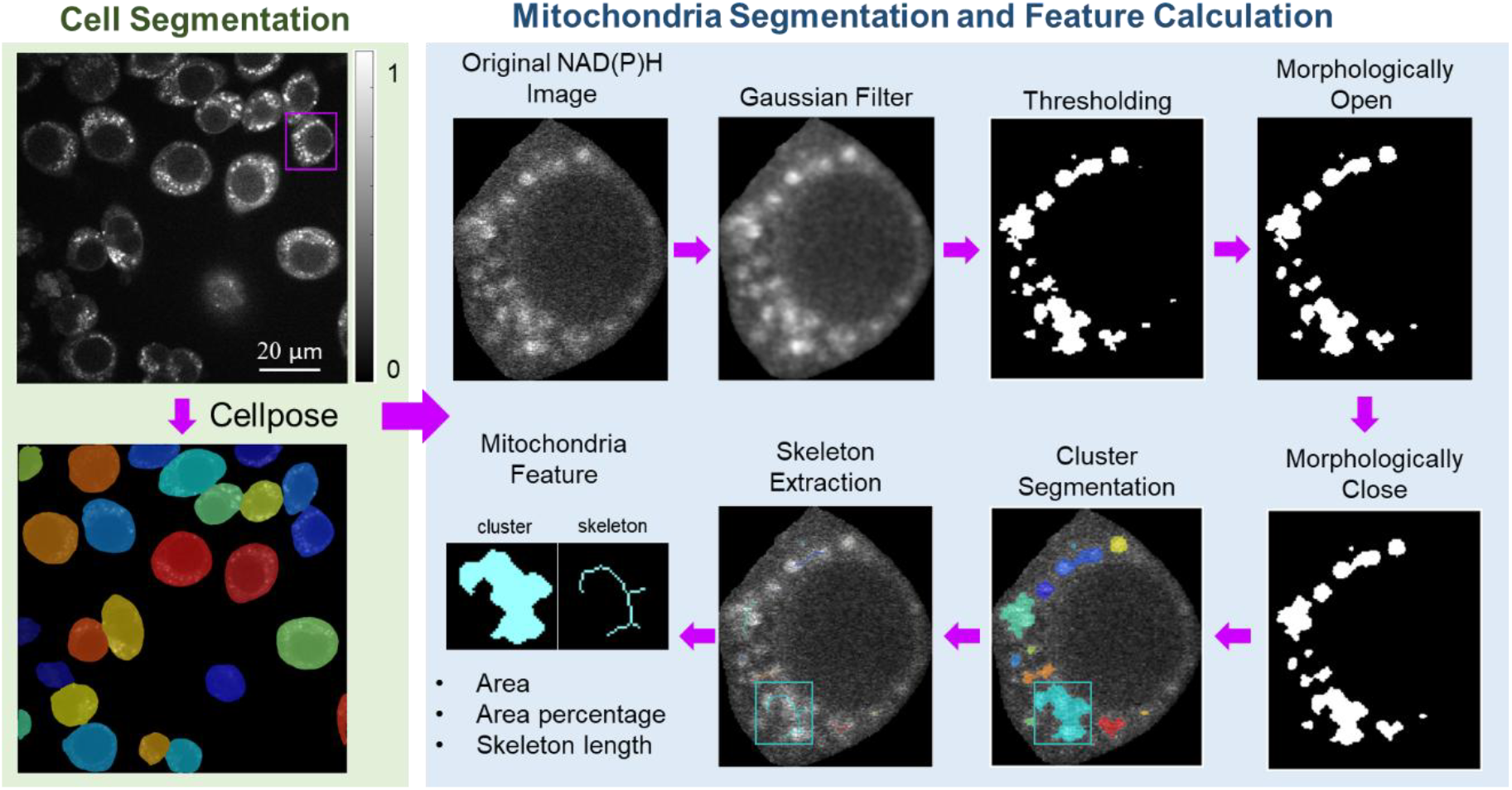
Schematic of mitochondria segmentation and feature calculation. Each NAD(P)H fluorescence image was segmented into individual cells using Cellpose. Then the mitochondria regions within each cell were segmented through filtering, thresholding, and morphologically open, and close. Clusters of mitochondria and associated skeletons were used to calculate the mitochondria features.

However, a single thresholding approach can lead to the misidentification of tiny bright regions (1 – 2 pixels) due to artifacts and noise after the Gaussian filter. To address this, morphological opening was performed. Firstly, the pre-identified mitochondrial mask was eroded and then dilated using a disk-shaped structuring element with a radius of 1 pixel. However, morphological opening can potentially break the connection between mitochondria objects, and create small holes within mitochondria clusters. To mitigate this effect, a morphological closing operation was applied. Using a disk-shaped structuring element with a radius of 2 pixels, the image was dilated and then eroded to obtain the final mitochondrial mask.

Different mitochondria clusters were segmented by an 8-connective criterion, which defines that two adjoining pixels belong to the same object if they are both on and connected along the horizontal, vertical, or diagonal direction. The skeleton of each cluster was extracted by eroding all objects to centerlines without altering the essential structure of the objects, using the MATLAB bwskel function. Finally, the area of the mitochondria clusters and the length of their skeletons were calculated to quantify the mitochondria features within each cell. The area percentage of the mitochondria within each cell was determined by calculating the ratio of the mitochondria area to the cellular area.

### Statistical analysis

Data analysis was performed in RStudio. A two-sided t-test was used to assess differences between WT and POLG BMDMs for each fluorescence lifetime variable and mitochondria feature. A significance value of 0.05 was chosen to indicate the presence of a significant difference. The Uniform Manifold Approximation and Projection (UMAP), an unsupervised machine learning model, was used for dimension reduction to visualize clustering within the autofluorescence imaging datasets [32]. The heterogeneity of NAD(P)H *τ*_*m*_, FAD *τ*_*m*_, and the redox ratio was analyzed by fitting their distributions with two Gaussians. Subsequently, the median values (*μ*_*1*_, *μ*_*2*_) and standard deviations (*σ*_*1*_, *σ*_*2*_) of these two Gaussians were used to calculate the bimodality index (BI) [33]:

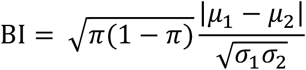

where 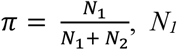 and *N*_*2*_ represent the number of cells in subpopulations. As previously defined, a BI > 1.1 indicates the presence of bimodality in the distribution [34].

## Results

### 3.1 Metabolic differences between WT and POLG macrophages

In autofluorescence images, the control BMDMs appear as circular shapes with nuclei darker than the cytoplasm due to lower concentrations of NAD(P)H and FAD in the nucleus (Fig. 2(a)). Activation of macrophages through TLR4 by LPS leads to increased cellular volume, cytoplasmic enlargement, and cell spreading (Fig. 2(a)).

**Figure 2.**
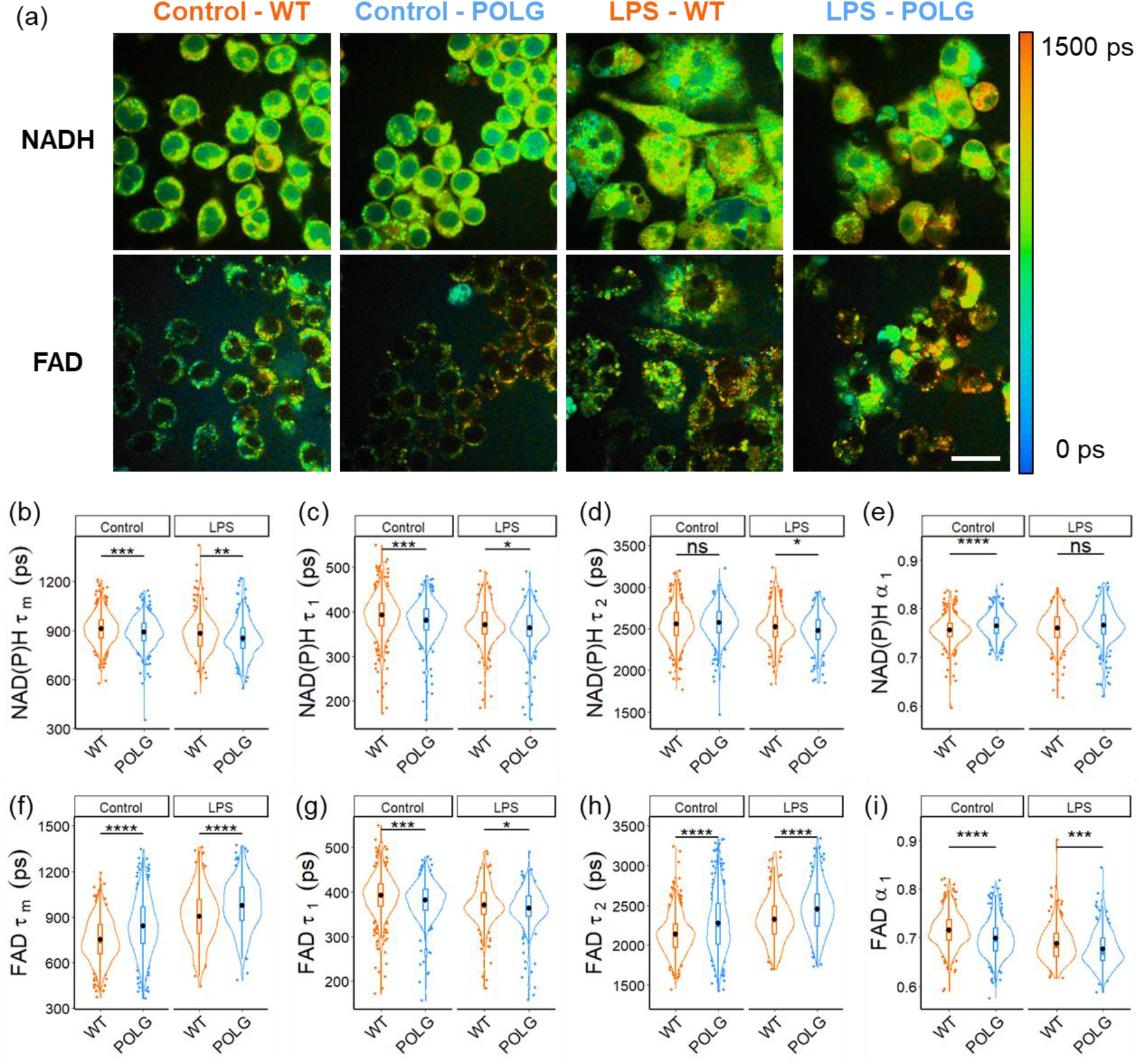
Autofluorescence lifetime imaging reveals different metabolic states between WT and POLG macrophages and due to LPS activation. (a) Representative NAD(P)H and FAD *τ*_*m*_ images of WT control and POLG macrophages with and without LPS treatment, scale bar = 25 *μm*. (b) NAD(P)H *τ*_*m*_ (c) NAD(P)H *τ*_*1*_ (d) NAD(P)H *τ*_*2*_ (e) NAD(P)H *α*_*1*_ (f) FAD *τ*_*m*_ (g) FAD *τ*_*1*_ (h) FAD *τ*_*2*_ and (i) FAD *α*_*1*_ of control and LPS stimulated WT and POLG BMDMs. *p < 0.05, **p < 0.01, ***p < 0.001, ****p < 0.0001 for two-sided student t-test. Statistics shown on plots are for WT-POLG comparisons within control or LPS treatment. Statistics for control-LPS comparisons for WT and POLG BMDMs are provided in Supplementary Table 2.

Autofluorescence lifetime imaging reveals differences in the NAD(P)H fluorescence lifetimes of POLG versus WT BMDMs and due to LPS treatment. POLG macrophages have a shorter free NAD(P)H lifetime (*τ*_*1*_), and a higher fraction of free NAD(P)H (*α*_*1*_), resulting in a shorter mean NAD(P)H lifetime (*τ*_*m*_) compared to the WT macrophages (Fig. 2(b)(c)(e)). The differences in short and average NAD(P)H lifetimes (*τ*_*1*_, *τ*_*m*_) between WT and POLG macrophages persist upon LPS activation, albeit to a lesser extent. Additionally, LPS treatment leads to a shorter mean NAD(P)H lifetime (*τ*_*m*_) in both WT and POLG macrophages (Sup. Table 2).

Similarly, FAD fluorescence lifetimes are different between WT and POLG BMDMs and due to LPS treatment. As compared with WT, POLG macrophages exhibit a shorter bound FAD lifetime (*τ*_*1*_, Fig. 2(g)), a longer free FAD lifetime (*τ*_*2*_, Fig. 2(h)), and a reduced fraction of bound FAD (*α*_*1*_, Fig. 2(i)). Consequently, POLG macrophages have an overall longer average FAD lifetime (*τ*_*m*_) than WT macrophages (Fig. 2(f)). In both WT and POLG macrophages upon LPS activation, both free and bound FAD lifetimes (*τ*_*1*_, *τ*_*2*_) increase, while the fraction (*α*_*1*_) of bound FAD decreases (Fig. 2(g)(h)(i), Sup. Table 2). These variations result in an overall increase in the average FAD lifetime (*τ*_*m*_) (Fig. 2(f), Sup. Table 2). The POLG BMDMs exhibited a lower redox ratio compared to WT BMDMs, and LPS activation eliminated these differences (Sup. Fig. 1). Furthermore, LPS stimulation leads to a significant increase in the redox ratio in both WT and POLG BMDMs as compared with the respective unstimulated WT or POLG BMDMs (Sup. Fig. 1(b), Sup. Table 2).

### 3.2 OMI resolves metabolic heterogeneity within WT and POLG BMDMs

Metabolic heterogeneity within the macrophage populations was analyzed by histogram assessment and calculation of the bimodality index (BI). The distribution of average NAD(P)H lifetime (*τ*_*m*_), average FAD lifetime (*τ*_*m*_), and redox ratio values for each macrophage group (WT control, WT LPS, POLG control, POLG LPS) were fit to two Gaussians. The BI was then calculated from the bimodal Gaussian fit and used to assess the presence of metabolic diversity within the cell population. LPS stimulated an increase in the BI to a value exceeding 1.1 for NAD(P)H *τ*_*m*_ and the optical redox ratio of the POLG macrophage population across two experimental replicates (Fig. 3(a)(c)(d)). Additionally, FAD *τ*_*m*_ demonstrates a bimodal distribution in the response of WT macrophages to LPS activation, with BI values exceeding 1.1. When calculating the BI from the combined data of three replicated experiments, a more pronounced bimodal separation of cell populations, characterized by higher BI values in NAD(P)H *τ*_*m*_, FAD *τ*_*m*_, and optical redox ratio, is observed in POLG BMDMs compared to WT BMDMs both under normal conditions and after LPS activation (Sup. Fig 2(a)). Moreover, LPS activation increases the BI of all lifetime variables, except for NAD(P)H *τ*_*m*_ in control WT BMDMs, for both WT and POLG BMDMs (Sup. Fig 2(b)).

**Figure 3.**
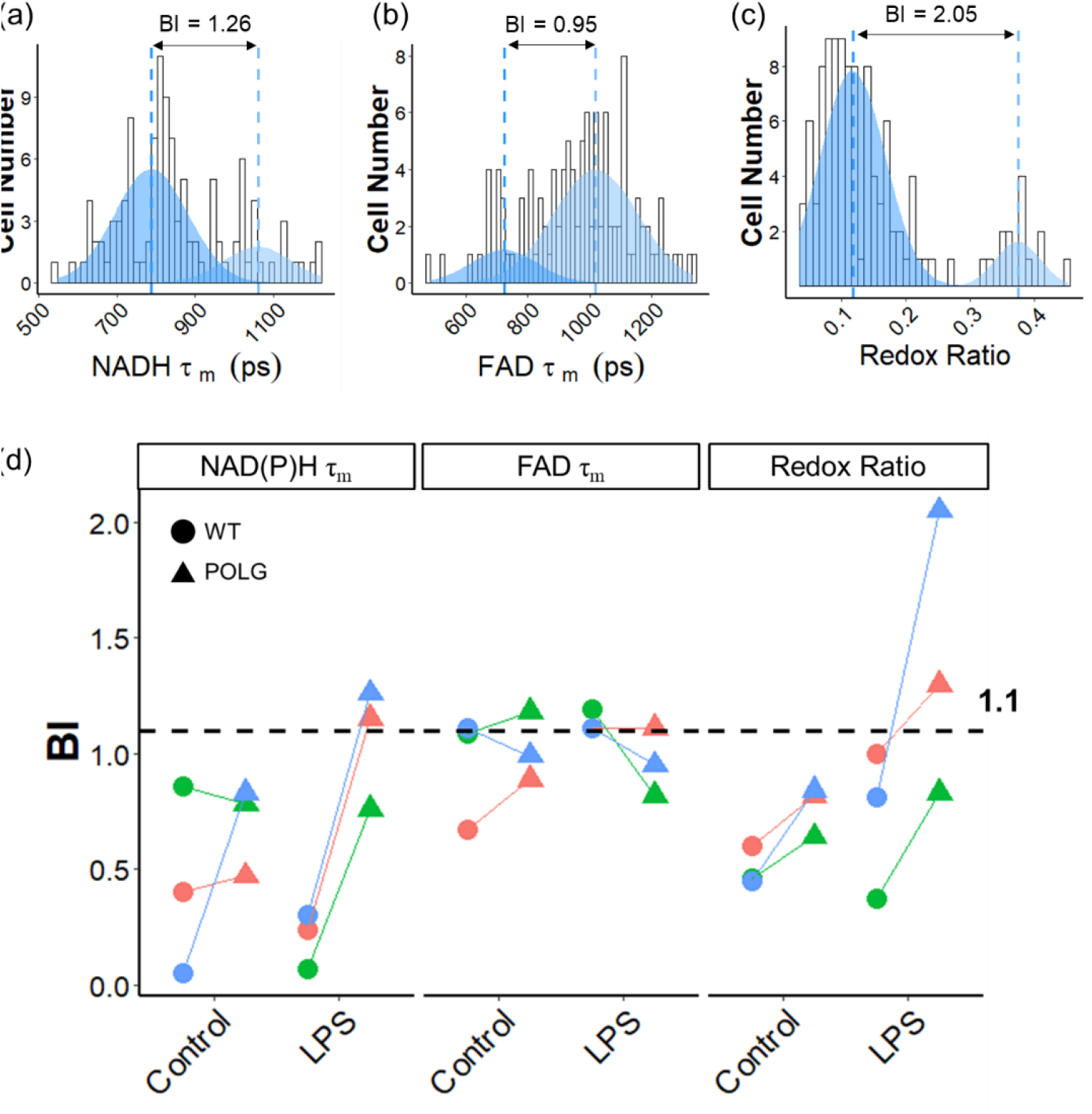
Metabolic heterogeneity analysis in WT and POLG macrophages using bimodal Gaussian fit. Distributions of the (a) mean NAD(P)H fluorescence lifetime *τ*_*m*_, (b) mean FAD fluorescence lifetime *τ*_*m*_, and (c) optical redox ratio (FAD/(FAD+NAD(P)H)) for LPS-treated POLG BMDMs, and the corresponding bi-modal Gaussian fit. (d) BI of NAD(P)H *τ*_*m*_, FAD *τ*_*m*_, and optical redox ratio of control, and LPS-treated WT and POLG BMDMs. Different colors represent different experimental replicates.

The analysis of the distribution of NADH *τ*_*m*_, FAD *τ*_*m*_, and redox ratio in the entire population shows that NADH *τ*_*m*_ and FAD *τ*_*m*_ exhibit lower skewness and kurtosis compared to the optical redox ratio (Sup. Fig. 3, Sup. Table 6). LPS treatment results in a higher standard deviation of NADH *τ*_*m*_ for both WT and POLG macrophages (Sup. Table 6).

### 3.3 OMI resolves intracellular heterogeneity within WT and POLG BMDMs

The intra-cellular standard deviation was calculated across the pixels of the cytoplasm area within each cell to evaluate intra-cellular heterogeneity. The intracellular standard deviation of NAD(P)H *τ*_*m*_ and the optical redox ratio are lower in POLG macrophages compared to WT macrophages in both control and LPS-treated BMDMs (Fig. 4(a)(c)). However, POLG BMDMs display a higher intra-cellular standard deviation of average FAD lifetime (*τ*_*m*_) than WT macrophages in both control and LPS treatment groups (Fig. 4(b)). Furthermore, LPS treatment does not significantly affect the standard deviation of NAD(P)H *τ*_*m*_ or FAD *τ*_*m*_ in either WT or POLG BMDMs (Sup. Fig. 4(a)(b)). However, LPS treatment increases the standard deviation of the optical redox ratio in both WT and POLG BMDMs (Sup. Fig. 4(c)).

**Figure 4.**
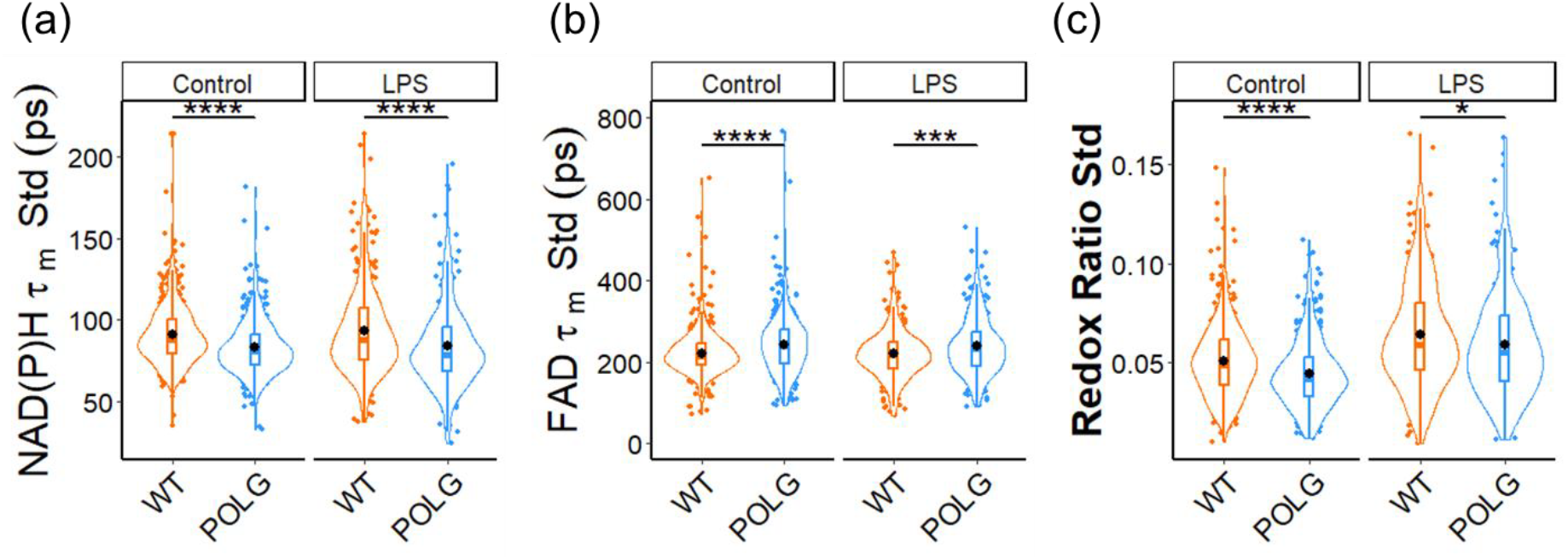
Intracellular heterogeneity within WT and POLG BMDMs. (a) Intra-cellular standard deviation of (a) mean NAD(P)H fluorescence lifetime *τ*_*m*_, (b) mean FAD fluorescence lifetime *τ*_*m*_, and (c) optical redox ratio for control and LPS-treated WT and POLG BMDMs. *p < 0.05, ***p < 0.001, and ****p < 0.0001 for two-sided student t-test.

### 3.3 Mitochondria differences between POLG and WT macrophages

The mitochondria exhibit brighter areas in the NAD(P)H fluorescence images due to the higher level of NAD(P)H within these organelles compared to the cytosol. This observation is validated by imaging cells stained with MitoTracker Deep Red, which specifically labels mitochondria with a fluorophore that has a distinct spectrum from NAD(P)H fluorescence (Fig. 5(a)). The brighter pixels in the NAD(P)H image correspond to the mitochondria within the cells (Fig. 5(a)), and the MATLAB mitochondrial segmentation pipeline identifies and segments the mitochondria regions from the NAD(P)H image, allowing for further mitochondria feature extraction (Fig. 5(a)).

**Figure 5.**
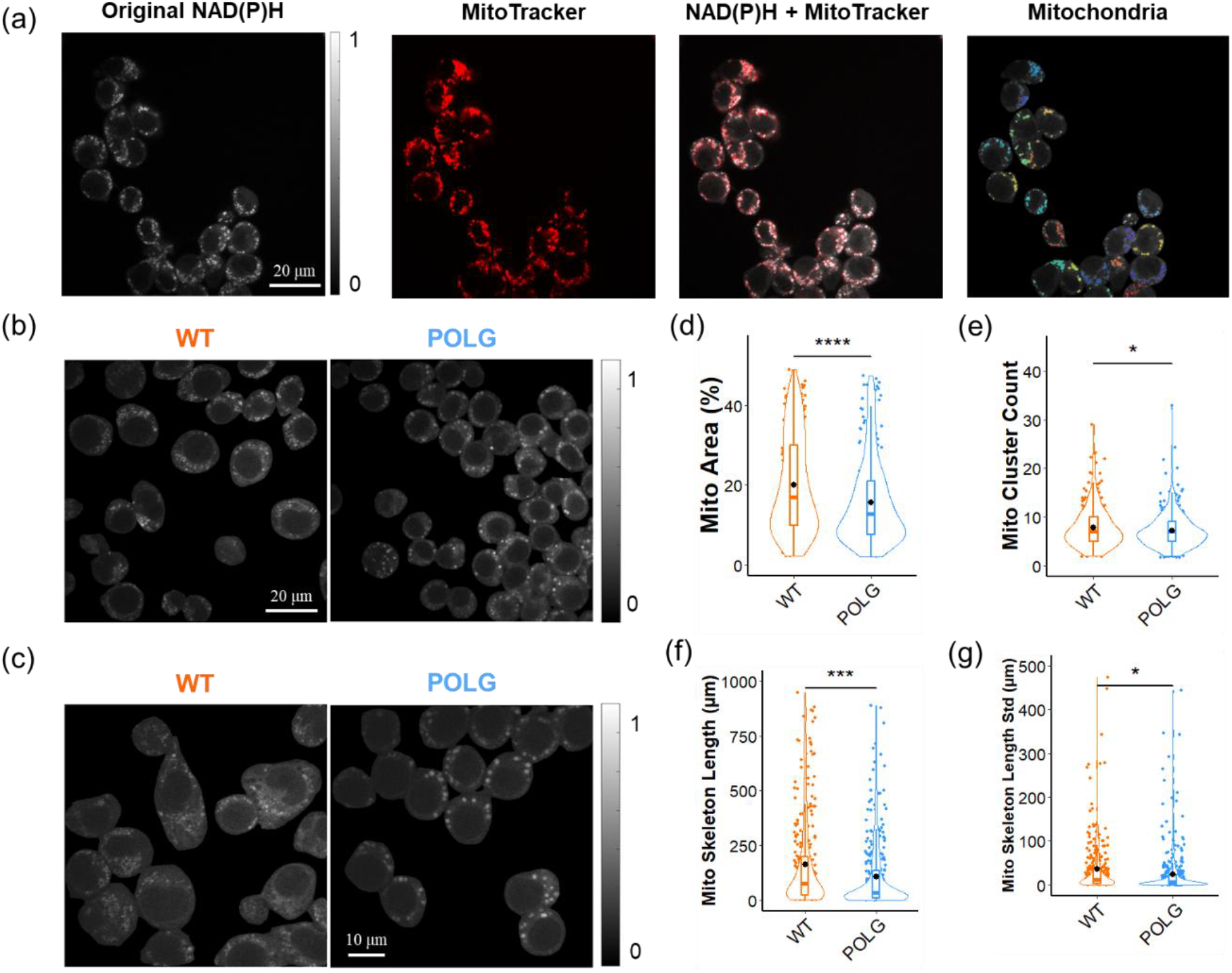
Assessment of mitochondria differences in control WT and POLG BMDMs. (a) Representative NAD(P)H, MitoTracker captured by a 100x (1.46 NA) objective, and the corresponding segmented mitochondria images for control WT BMDMs. (b) Representative NAD(P)H autofluorescence images of WT and POLG macrophages captured by a 100x (1.46 NA) objective (c) Representative NAD(P)H autofluorescence images of WT and POLG macrophages captured by a 150x (1.35 NA) objective (d) mitochondria area percentage over whole cell area (e) Number of mitochondria clusters (f) Sum of length of mitochondria skeleton (g) Standard deviation of length of mitochondria skeleton in WT and POLG macrophages. *p < 0.05, ***p < 0.001, ****p < 0.0001 for two-sided student t-test.

The distinct characteristics of mitochondria between POLG and WT BMDMs can be observed from NAD(P)H intensity images (Fig. 5(b)(c)). In WT BMDMs, mitochondria form larger clusters and exhibit a higher overall abundance compared to POLG BMDMs (Fig. 5(b)), as illustrated in images captured with a 150x objective (Fig. 5(c)). Quantitative analysis of the mitochondria in the NAD(P)H images reveals that mitochondria within WT BMDMs comprise a higher percentage of the whole cell area and a greater number of mitochondria clusters (Fig. 4(d)(e)). Examination of the length of the skeleton of mitochondria clusters indicates that WT BMDMs exhibit longer skeleton lengths and greater standard deviations than POLG BMDMs within each cell (Fig. 5(f)(g)). Additionally, the NAD(P)H images captured by the 150x objective allow for quantitative discrimination of mitochondria morphology between WT and POLG BMDMs, with results consistent with those obtained from analyzing images acquired with the 100x objective (Sup. Fig. 5). Furthermore, the mitochondria within LPS-treated WT BMDMs also exhibit a higher percentage of the whole cell area and a longer skeleton length than the POLG BMDMs (Sup Fig. 6).

### 3.4 Metabolic drift in POLG BMDMs over generations

The significant metabolic differences, as revealed by OMI, between WT and POLG BMDMs remain consistent across multiple generations of BMDMs (Fig. 2). However, a closer examination using 3D scatter plots of three representative OMI endpoints, NAD(P)H *τ*_*m*_, FAD *τ*_*m*_, and optical redox ratio show increased overlapping areas of the WT and POLG cells at higher generations (Fig. 6(a)(b)). Notably, the difference in the NAD(P)H *τ*_*m*_ between WT and POLG BMDMs becomes less pronounced in higher generations of cells [29] (Fig. 2(b), Sup. Fig. 7(a)), however, the difference in the FAD *τ*_*m*_ becomes more significant (Fig. 2(f), Sup. Fig. 7(b)). Furthermore, we used the autofluorescence lifetime features including NAD(P)H *τ*_*m*_, *τ*_*1*_, *τ*_*2*_, *α*_*1*_; FAD *τ*_*m*,_ *τ*_*1*_, *τ*_*2*_, *α*_*1*_; and optical redox ratio, for a 2D projection using UMAP. For the younger generation of cells, WT BMDMs exhibit a separate cluster from the POLG BMDMs (Fig. 6(c)). However, the separation between the groups is reduced and an increased amount of overlap between WT and POLG BMDMs is be observed for the UMAP generated from the optical metabolic imaging features of the older cell generation (Fig. 6(d)).

**Figure 6.**
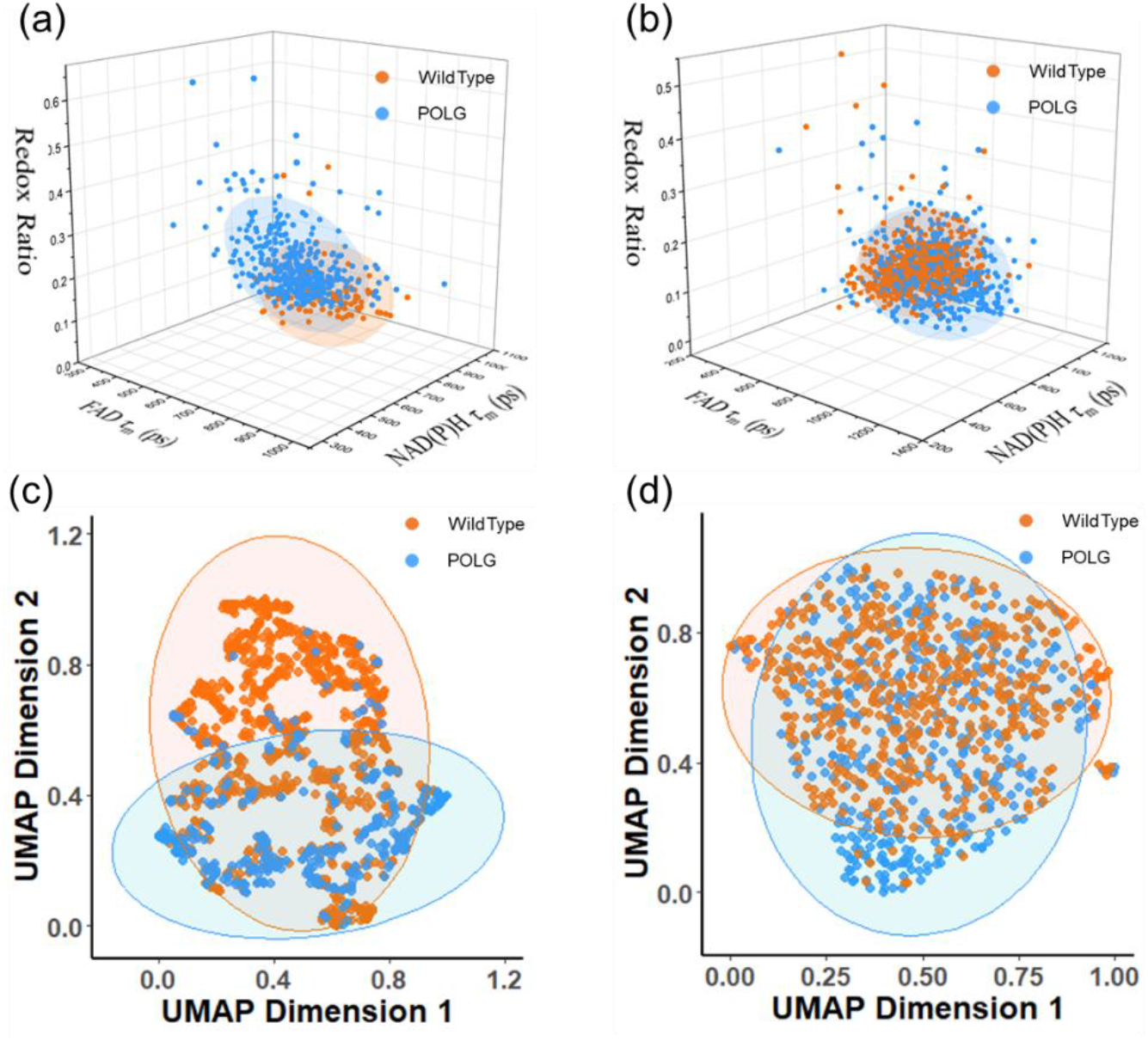
Convergence of metabolic profiles between WT and POLG BMDMs over generations. Scatter plot of NAD(P)H *τ*_*m*_, FAD *τ*_*m*_, and redox ratio for WT and POLG BMDMs of (a) younger (7^th^ - 10^th^) generation and (b) older (35^th^ – 40^th^) generation, the ellipsoid represents 85% of data coverage. UMAP representations of autofluorescence lifetime features (NAD(P)H *τ*_*m*_, *τ*_*1*_, *τ*_*2*_, *α*_*1*_; FAD *τ*_*m*_, *τ*_*1*_, *τ*_*2*_, *α*_*1*_; and redox ratio) for WT and POLG BMDMs of (c) younger (7^th^ -10^th^) generation and (d) older (35^th^ – 40^th^) generation, the ellipsoid represents 95% of the group.

## Discussion

POLG mutations lead to mitochondria disorders resulting in various diseases such as Alpers-Huttenlocher syndrome, childhood myocerebrohepatopathy spectrum, and myoclonic epilepsy myopathy sensory ataxia. Recently, several mouse models of POLG-related diseases have been developed, with the POLG mutator mouse being the most well-studied model [35, 36]. Previous studies have shown the inflammation contributes to increased oxidative stress and aerobic glycolysis in a population of BMDMs from POLG mutator mice [6]. Furthermore, POLG mutations can be heterogeneous at the mitochondria level, contributing to a wide range of clinical manifestations in individuals [5]. Hence, developing a non-invasive technology that can track metabolic shifts and characterize mitochondria function during immune cell activation could significantly contribute to studies of the effects of POLG mutations and provide a testbed for potential therapies. Optical imaging of NAD(P)H and FAD is sensitive to metabolic differences between WT and POLG BMDMs [29] and can resolve subcellular structures when using a high magnification objective. In this paper, we illustrate the capabilities and potential of OMI to measure the metabolic variations of POLG BMDMs and provide a customized image process technique to quantify differences in mitochondria between WT and POLG BMDMs.

Autofluorescence lifetime imaging of NAD(P)H and FAD identifies the metabolic variations of WT and POLG BMDMs. The shorter lifetime of NAD(P)H, a higher fraction of free NAD(P)H, and lower optical redox ratio values in POLG BMDMs suggests a more glycolytic status compared to WT BMDMs (Fig. 2(b-e)). This observation aligns with findings from Seahorse metabolic assay experiments conducted previously [29]. Additionally, the increase in the average FAD fluorescence lifetime (*τ*_*m*_) in POLG cells, attributed to a glycolytic phenotype, has also been observed in breast cancer cells under OXPHOS inhibition using cyanide treatment [18]. LPS activates various transcription factors, signal transducers, activators of transcription molecules, and an enhanced flux through glycolysis and fatty acid synthesis [37]. OMI reveals a significant decrease in average NAD(P)H lifetime (*τ*_*m*_) and an increase in average FAD lifetime (*τ*_*m*_) in both WT and POLG macrophages after LPS treatment (Sup. Table 2), suggesting metabolic variations towards greater dependence on glycolysis. Even though the macrophages here were not further polarized to proinflammatory phenotypes (M1) by interferons gamma (IFNγ), variations in the NAD(P)H and FAD lifetime remained consistent with what has been observed in M1 macrophages [38]. Furthermore, the minimal differences in NAD(P)H lifetime (*τ*_*m*_) and the significant overlaps in OMI feature projection between WT and POLG BMDMs of older generations suggest a convergence of metabolic profiles over passages (Fig. 6). This convergence could be due to epigenetic changes and environmental influences of multiple cell passages. While further research is needed to confirm this hypothesis, this observation should motivate researchers to closely monitor passage number as a potential confounding factor of POLG mutator BMDMs. The OMI technique holds the potential to noninvasively track cell differences over generations.

Fluorescence lifetime measurements are more robust than intensity measurements. Fluorescence lifetime measurements are self-referenced and are less susceptible to artifacts arising from scattered light, non-uniform illumination, inner filter effects, and detector gain as compared to intensity imaging [39, 40]. Herein, the BMDMs exhibit a more uniform distribution of NAD(P)H and FAD lifetime *τ*_*m*_ than the redox ratio with a lower skewness and kurtosis for the cell population (Sup. Fig. 3, Sup. Table 6). Interestingly, the redox ratio increases for both WT and POLG after LPS treatment (Sup. Fig. 1), which was also observed in a previous study showing an increase in redox ratio in M1 macrophages after 24 hours [41]. The increase in the redox ratio is due to a higher level of FAD, suggesting a higher reduced status, probably from various signal pathways triggered by LPS challenging, or additional fluorescent flavin molecules generated by the LPS challenge [42].

OMI detects metabolism at a single cell level, allowing for the identification of subpopulation cells within unknown, heterogeneous populations when incorporated with cell segmentation. Previously, subpopulation analysis of OMI indexes has been applied to discriminate intracellular heterogeneity among cell types [7, 43] and quantify heterogeneous responses to chemotherapy [33]. In this study, we utilized OMI to measure cellular metabolism and assess the heterogeneity in BMDM populations from different groups to better understand the influence of POLG mutations. The immunological challenge with LPS leads to metabolic subpopulations with a higher BI compared to control groups, especially for POLG BMDMs, which have a BI above 1.1 for NAD(P)H *τ*_*m*_ and redox ratio (Fig. 3(d)), suggesting heterogeneous immune responses of BMDMs. Combining datasets from each replicate reduces the BI values, as it compensates for imbalances and disparities in metabolic states and immune responses. Nonetheless, higher BI values are observed in POLG mutation BMDMs than in WT BMDMs (Sup. Fig. 2(a)), implicating the heterogeneous effects of the POLG mutation. The bimodal basal metabolism and immune responses in POLG-mutation BMDMs highlight the clinical challenges associated with POLG-related disease heterogeneity in affected patients and unpredictable disease courses [1].

OMI allows for the quantification of mitochondrial differences between WT and POLG BMDMs. As mitochondria contain a higher level of NAD(P)H than the cytosol [19], the mitochondria regions appear brighter in NAD(P)H autofluorescent images, allowing them to be segmented based on a threshold method, and pinpointing the autofluorescence lifetime features in the mitochondrial regions [44-46]. Herein, the MitoTracker image was captured sequentially after the NAD(P)H image, with about a 10-second delay in a different channel. Considering the dynamic movements of mitochondria, and potential mismatches between images in different channels, the mitochondrial regions can be identified as the brighter pixels in the NAD(P)H image, based on the staining patterns in the MitoTracker image. A customized image processing pipeline in MATLAB allows segmentation of mitochondria regions from NAD(P)H images, enabling the quantification of key mitochondria differences between WT and POLG BMDMs. A higher percentage of mitochondria area and longer skeleton length are observed in WT BMDMs than in POLG BMDMs (Fig. 5(d)(f)). This is likely due to disruptions in the replication and repair of the mitochondrial genome from POLG mutations, resulting in variations in mitochondria network structures. Additional morphological features, such as circularity, orientation, and perimeter can be obtained by further analyzing the segmented mitochondrial regions. The LPS-treated WT BMDMs exhibit a higher percentage of mitochondria area than POLG BMDMs (Sup. Fig. 6(b)(d)). However, LPS treatment induces mitochondrial fission and fragmentation [47], resulting in more concentrated mitochondrial clusters, as observed in the NAD(P)H images (Sup. Fig. 6(a)). This creates additional challenges in segmenting individual mitochondrial components and precisely quantifying their characteristics. Moreover, intracellular standard deviations in POLG BMDMs are lower than WT BMDMs for NAD(P)H *τ*_*m*_ and redox ratio (Fig. 4(a)(c)), but higher for FAD *τ*_*m*_ (Fig. 4(b)), suggesting different mitochondria morphological patterns within cells between WT and POLG BMDMs.

## Conclusion

In this paper, we demonstrate the capability of OMI to detect metabolic differences between WT and POLG BMDMs, as well as identify subpopulations of responses under immunological activation. Furthermore, we propose an image processing pipeline to segment mitochondrial regions from NAD(P)H images and extract mitochondria morphology metrics including percent area and skeleton length. The WT BMDMs exhibit a higher percentage of mitochondria regions with longer skeleton lengths compared to POLG BMDMs. OMI provides a label-free imaging technique that reveals metabolic activity while resolving intracellular information at the single-cell level. This is advantageous for dynamic studies aimed at detecting metabolism and tracking variations within individual cells and across cell populations. Furthermore, as POLG mutations are associated with various mitochondrial diseases, the development of noninvasive imaging tools to assess metabolic status and mitochondrial structures in POLG mutator cells could provide critical insights into their heterogeneity and responses to therapy.

## Disclosures

The authors have no relevant financial interests in this manuscript and no potential conflicts of interest.

## Acknowledgments

The authors would like to thank Dr. Phillip West, and Dr. Yuanjiu Lei for providing the wild-type and POLG mutator bone marrow-derived macrophages and their valuable suggestions. A.J.W. was supported by NIH NIGMS R35 GM1412990.

## Notes

### Competing Interest Statement

The authors have declared no competing interest.

## References

1. Rahman, S. and W.C. Copeland, POLG-related disorders and their neurological manifestations. Nat Rev Neurol, 2019. 15(1): p. 40–52.

2. Young, M.J. and W.C. Copeland, Human mitochondrial DNA replication machinery and disease. Curr Opin Genet Dev, 2016. 38: p. 52–62.

3. Singh, B., et al., Mitochondrial DNA Polymerase POLG1 Disease Mutations and Germline Variants Promote Tumorigenic Properties. PLOS ONE, 2015. 10(10): p. e0139846.

4. Copeland, W.C., The mitochondrial DNA polymerase in health and disease. Subcell Biochem, 2010. 50: p. 211–22.

5. Cakmak, C. and H. Zempel, A perspective on human cell models for POLG-spectrum disorders: advantages and disadvantages of CRISPR-Cas-based vs. patient-derived iPSC models. Medizinische Genetik, 2021. 33(3): p. 245–249.

6. Lei, Y., et al., Elevated type I interferon responses potentiate metabolic dysfunction, inflammation, and accelerated aging in mtDNA mutator mice. Science Advances, 2021. 7(22): p. eabe7548.

7. Walsh, A.J. and M.C. Skala, Optical metabolic imaging quantifies heterogeneous cell populations. Biomed Opt Express, 2015. 6(2): p. 559–73.

8. Georgakoudi, I. and K.P. Quinn, Optical imaging using endogenous contrast to assess metabolic state. Annu Rev Biomed Eng, 2012. 14: p. 351–67.

9. Chance, B., et al., Oxidation-reduction ratio studies of mitochondria in freeze-trapped samples. Journal of Biological Chemistry, 1979. 254(11): p. 4764–4771.

10. Kolenc, O.I. and K.P. Quinn, Evaluating Cell Metabolism Through Autofluorescence Imaging of NAD(P)H and FAD. Antioxid Redox Signal, 2019. 30(6): p. 875–889.

11. Varone, A., et al., Endogenous two-photon fluorescence imaging elucidates metabolic changes related to enhanced glycolysis and glutamine consumption in precancerous epithelial tissues. Cancer Res, 2014. 74(11): p. 3067–75.

12. Huang, S., A.A. Heikal, and W.W. Webb, Two-photon fluorescence spectroscopy and microscopy of NAD(P)H and flavoprotein. Biophys J, 2002. 82(5): p. 2811–25.

13. Lakowicz, J.R., et al., Fluorescence lifetime imaging of free and protein-bound NADH. Proc Natl Acad Sci U S A, 1992. 89(4): p. 1271–5.

14. Nakashima, N., et al., Picosecond fluorescence lifetime of the coenzyme of D-amino acid oxidase. J Biol Chem, 1980. 255(11): p. 5261–3.

15. Melissa C. Skala, et al., In vivo multiphoton microscopy of NADH and FAD redox states, fluorescence lifetimes, and cellular morphology in precancerous epithelia. PNAS, 2007. 104(49): p. 19494–19499.

16. Walsh, A.J., et al., Classification of T-cell activation via autofluorescence lifetime imaging. Nat Biomed Eng, 2021. 5(1): p. 77–88.

17. Qian, T., et al., Label-free imaging for quality control of cardiomyocyte differentiation. Nat Commun, 2021. 12(1): p. 4580.

18. Walsh, A.J., et al., Optical metabolic imaging identifies glycolytic levels, subtypes, and early-treatment response in breast cancer. Cancer Res, 2013. 73(20): p. 6164–74.

19. Stein, L.R. and S. Imai, The dynamic regulation of NAD metabolism in mitochondria. Trends Endocrinol Metab, 2012. 23(9): p. 420–8.

20. Jonathan M Levitt, et al., Diagnostic cellular organization features extracted from autofluorescence images. Opt Lett., 2007. 32(22): p. 3305 – 7.

21. Xylas, J., et al., Noninvasive assessment of mitochondrial organization in three-dimensional tissues reveals changes associated with cancer development. Int J Cancer, 2015. 136(2): p. 322–32.

22. Hu, L., et al., Label-free spatially maintained measurements of metabolic phenotypes in cells. Front Bioeng Biotechnol, 2023. 11: p. 1293268.

23. Wang, Z.J., et al., Classifying T cell activity in autofluorescence intensity images with convolutional neural networks. J Biophotonics, 2020. 13(3): p. e201960050.

24. Ramey, N.A., et al., Imaging mitochondria in living corneal endothelial cells using autofluorescence microscopy. Photochem Photobiol, 2007. 83(6): p. 1325–9.

25. Allen, C.H., et al., Label-free two-photon imaging of mitochondrial activity in murine macrophages stimulated with bacterial and viral ligands. Sci Rep, 2021. 11(1): p. 14081.

26. Lefebvre, A., et al., Automated segmentation and tracking of mitochondria in live-cell time-lapse images. Nat Methods, 2021. 18(9): p. 1091–1102.

27. Valente, A.J., et al., A simple ImageJ macro tool for analyzing mitochondrial network morphology in mammalian cell culture. Acta Histochem, 2017. 119(3): p. 315–326.

28. Chaudhry, A., R. Shi, and D.S. Luciani, A pipeline for multidimensional confocal analysis of mitochondrial morphology, function, and dynamics in pancreatic β-cells. American Journal of Physiology-Endocrinology and Metabolism, 2020. 318(2): p. E87–E101.

29. Hu, L., et al., 3D convolutional neural networks predict cellular metabolic pathway use from fluorescence lifetime decay data. APL Bioeng, 2024. 8(1): p. 016112.

30. Stringer, C., et al., Cellpose: a generalist algorithm for cellular segmentation. Nat Methods, 2021. 18(1): p. 100–106.

31. Pachitariu, M. and C. Stringer, Cellpose 2.0: how to train your own model. Nat Methods, 2022. 19(12): p. 1634–1641.

32. McInnes, L., J. Healy, and J. Melville, Umap: Uniform manifold approximation and projection for dimension reduction. arXiv preprint 1802.03426, 2018.

33. Shirshin, E.A., et al., Label-free sensing of cells with fluorescence lifetime imaging: The quest for metabolic heterogeneity. Proceedings of the National Academy of Sciences, 2022. 119(9): p. e2118241119.

34. Wang, J., et al., The bimodality index: a criterion for discovering and ranking bimodal signatures from cancer gene expression profiling data. Cancer Inform, 2009. 7: p. 199–216.

35. Trifunovic, A., et al., Premature ageing in mice expressing defective mitochondrial DNA polymerase. Nature, 2004. 429(6990): p. 417–23.

36. Kujoth, G.C., et al., Mitochondrial DNA mutations, oxidative stress, and apoptosis in mammalian aging. Science, 2005. 309(5733): p. 481–4.

37. Marrocco, A. and L.A. Ortiz, Role of metabolic reprogramming in pro-inflammatory cytokine secretion from LPS or silica-activated macrophages. Front Immunol, 2022. 13: p. 936167.

38. Smokelin, I., et al., Optical changes in THP-1 macrophage metabolism in response to pro- and anti-inflammatory stimuli reported by label-free two-photon imaging. J Biomed Opt, 2020. 25(1): p. 1–14.

39. Becker, W., Fluorescence lifetime imaging--techniques and applications. J Microsc, 2012. 247(2): p. 119–36.

40. Datta, R., et al., Fluorescence lifetime imaging microscopy: fundamentals and advances in instrumentation, analysis, and applications. J Biomed Opt, 2020. 25(7): p. 1–43.

41. Bess, S.N., et al., Autofluorescence imaging of endogenous metabolic cofactors in response to cytokine stimulation of classically activated macrophages. Cancer Metab, 2023. 11(1): p. 22.

42. Al-Harbi, N.O., et al., Riboflavin attenuates lipopolysaccharide-induced lung injury in rats. Toxicol Mech Methods, 2015. 25(5): p. 417–23.

43. Chacko, J.V. and K.W. Eliceiri, Autofluorescence lifetime imaging of cellular metabolism: Sensitivity toward cell density, pH, intracellular, and intercellular heterogeneity. Cytometry A, 2019. 95(1): p. 56–69.

44. Yu, Q. and A.A. Heikal, Two-photon autofluorescence dynamics imaging reveals sensitivity of intracellular NADH concentration and conformation to cell physiology at the single-cell level. J Photochem Photobiol B, 2009. 95(1): p. 46–57.

45. Alam, S.R., et al., Investigation of Mitochondrial Metabolic Response to Doxorubicin in Prostate Cancer Cells: An NADH, FAD and Tryptophan FLIM Assay. Sci Rep, 2017. 7(1): p. 10451.

46. Hu, L., et al., Fluorescence intensity and lifetime redox ratios detect metabolic perturbations in T cells. Biomed Opt Express, 2020. 11(10): p. 5674–5688.

47. He, M., et al., Lipopolysaccharide induces human olfactory ensheathing glial apoptosis by promoting mitochondrial dysfunction and activating the JNK-Bnip3-Bax pathway. Cell Stress and Chaperones, 2019. 24(1): p. 91–104.

