## Supplementary Information for "Optical metabolic imaging identifies metabolic shifts and mitochondria heterogeneity in POLG mutator macrophages"

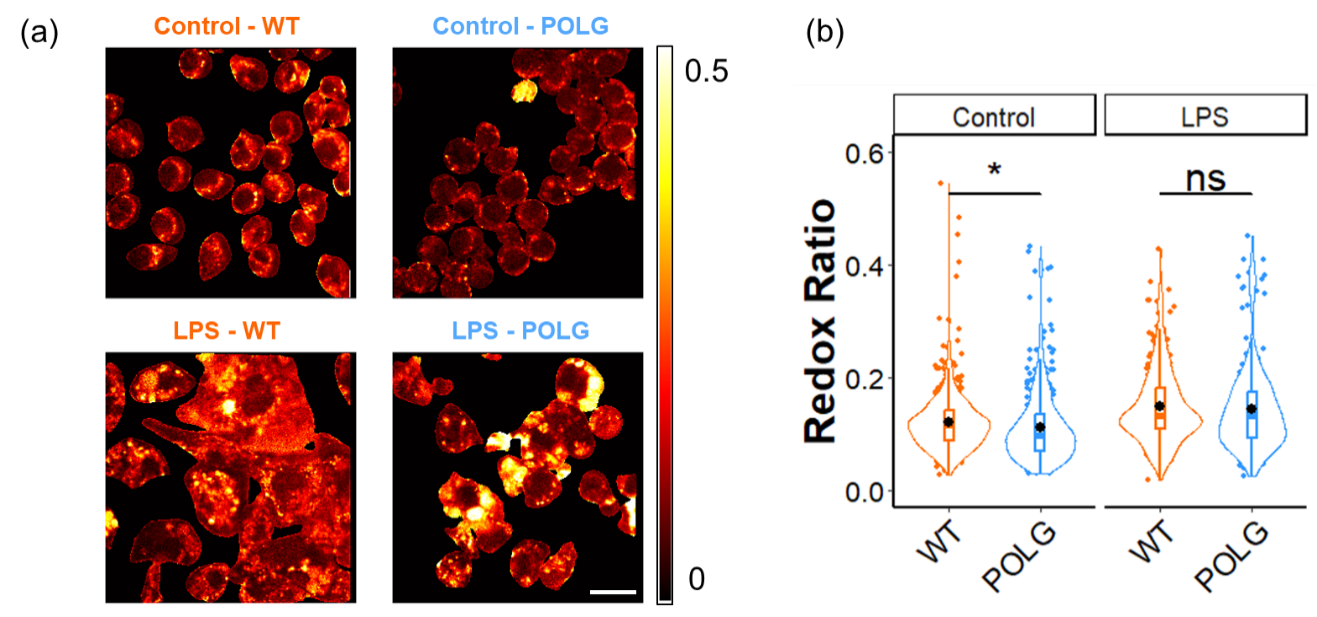

**Figure 1. Optical** redox ratio (FAD/(FAD+NAD(P)H)) variations of WT and POLG BMDMs with LPS treatment. (a) Representative redox ratio images of wild-type and POLG BMDMs with LPS treatment, scale bar = 20 *μm.* (b) The redox ratio of each cell for WT and POLG BMDMs with LPS treatment. * p < 0.05 for two-sided student t-test.

Table 1. Number of cells in each group

|  | Groups | Cell Number |
| --- | --- | --- |
| Lifetime Imaging | WT Control | 542 |
|  | POLG Control | 429 |
|  | WT LPS | 268 |
|  | POLG LPS | 248 |
| Intensity Imaging | WT Control (100X) | 304 |
|  | POLG Control (100X) | 337 |
|  | WT Control (150X) | 109 |
|  | POLG Control (150X) | 116 |

Table 2. Effects of LPS treatment on lifetime endpoints

|  | NAD(P)H | | | | FAD | | | | Redox Ratio |
| --- | --- | --- | --- | --- | --- | --- | --- | --- | --- |
|  | *α_1_* | *τ_1_* | *τ_2_* | *τ_m_* | *α_1_* | *τ_1_* | *τ_2_* | *τ_m_* |  |
| WT LPS | **-** | **↓** | **-** | **↓** | **↓** | **↑** | **↑** | **↑** | **↑** |
| Significant Level | **-** | ******** | **-** | ******* | ******** | ******** | ******** | ******** | ******** |
| POLG LPS | **-** | **↓** | **↓** | **↓** | **↓** | **↑** | **↑** | **↑** | **↑** |
| Significant Level | **-** | ******** | ******** | ******** | ******** | ******** | ******** | ******** | ******** |

Table 3. Bimodal Gaussian analysis of different BMDMs for the first experimental replicate

|  |  | N_1_ | N_2_ | μ_1_ | μ_2_ | σ_1_ | σ_2_ | BI |
| --- | --- | --- | --- | --- | --- | --- | --- | --- |
| Control WT | NADH τ_m_ | 64 | 157 | 851 | 926 | 102 | 71 | 0.40 |
|  | FAD τ_m_ | 2 | 219 | 1171 | 724 | 23 | 139 | 0.67 |
|  | Redox ratio | 211 | 10 | 0.11 | 0.32 | 0.035 | 0.142 | 0.60 |
| Control POLG | NADH τ_m_ | 10 | 68 | 865 | 907 | 11 | 75 | 0.47 |
|  | FAD τ_m_ | 74 | 4 | 765 | 1182 | 146 | 70 | 0.89 |
|  | Redox ratio | 69 | 9 | 0.17 | 0.24 | 0.034 | 0.027 | 0.82 |
| LPS WT | NADH τ_m_ | 10 | 51 | 806 | 833 | 20 | 89 | 0.24 |
|  | FAD τ_m_ | 7 | 54 | 627 | 902 | 34 | 186 | 1.11 |
|  | Redox ratio | 21 | 41 | 0.11 | 0.20 | 0.029 | 0.067 | 1.00 |
| LPS POLG | NADH τ_m_ | 8 | 57 | 1005 | 880 | 24 | 54 | 1.15 |
|  | FAD τ_m_ | 49 | 16 | 932 | 1108 | 145 | 33 | 1.11 |
|  | Redox ratio | 24 | 41 | 0.12 | 0.19 | 0.027 | 0.029 | 1.30 |

BI > 1.1 denoted with shading.

Table 4. Bimodal Gaussian analysis of different BMDMs for the second experimental replicate

|  |  | N_1_ | N_2_ | μ_1_ | μ_2_ | σ_1_ | σ_2_ | BI |
| --- | --- | --- | --- | --- | --- | --- | --- | --- |
| Control WT | NADH τ_m_ | 88 | 12 | 956 | 1129 | 83 | 50 | 0.86 |
|  | FAD τ_m_ | 67 | 34 | 754 | 945 | 92 | 75 | 1.09 |
|  | Redox ratio | 6 | 94 | 0.25 | 0.13 | 0.080 | 0.043 | 0.46 |
| Control POLG | NADH τ_m_ | 2 | 123 | 603 | 923 | 29 | 86 | 0.78 |
|  | FAD τ_m_ | 103 | 22 | 799 | 1038 | 99 | 59 | 1.18 |
|  | Redox ratio | 113 | 12 | 0.10 | 0.22 | 0.032 | 0.100 | 0.64 |
| LPS WT | NADH τ_m_ | 85 | 12 | 876 | 884 | 100 | 10 | 0.07 |
|  | FAD τ_m_ | 83 | 14 | 920 | 1144 | 97 | 45 | 1.19 |
|  | Redox ratio | 32 | 65 | 0.12 | 0.14 | 0.009 | 0.056 | 0.37 |
| LPS POLG | NADH τ_m_ | 22 | 50 | 747 | 851 | 57 | 71 | 0.76 |
|  | FAD τ_m_ | 23 | 49 | 873 | 1052 | 77 | 134 | 0.82 |
|  | Redox ratio | 2 | 70 | 0.24 | 0.12 | 0.014 | 0.036 | 0.83 |

BI > 1.1 denoted with shading.

Table 5. Bimodal Gaussian analysis of different BMDMs for the third experimental replicate

|  |  | N_1_ | N_2_ | μ_1_ | μ_2_ | σ_1_ | σ_2_ | BI |
| --- | --- | --- | --- | --- | --- | --- | --- | --- |
| Control WT | NADH τ_m_ | 212 | 9 | 888 | 863 | 74 | 157 | 0.05 |
|  | FAD τ_m_ | 126 | 95 | 666 | 851 | 80 | 86 | 1.11 |
|  | Redox ratio | 14 | 207 | 0.19 | 0.11 | 0.059 | 0.032 | 0.45 |
| Control POLG | NADH τ_m_ | 3 | 222 | 755 | 870 | 3 | 79 | 0.83 |
|  | FAD τ_m_ | 73 | 152 | 659 | 951 | 130 | 146 | 0.99 |
|  | Redox ratio | 213 | 12 | 0.08 | 0.29 | 0.028 | 0.105 | 0.84 |
| LPS WT | NADH τ_m_ | 70 | 39 | 893 | 965 | 76 | 180 | 0.30 |
|  | FAD τ_m_ | 5 | 104 | 1302 | 854 | 5 | 104 | 1.11 |
|  | Redox ratio | 82 | 27 | 0.12 | 0.23 | 0.042 | 0.082 | 0.81 |
| LPS POLG | NADH τ_m_ | 86 | 26 | 788 | 1060 | 94 | 88 | 1.26 |
|  | FAD τ_m_ | 21 | 91 | 724 | 1019 | 106 | 137 | 0.95 |
|  | Redox ratio | 98 | 14 | 0.12 | 0.37 | 0.050 | 0.035 | 2.05 |

BI > 1.1 denoted with shading.

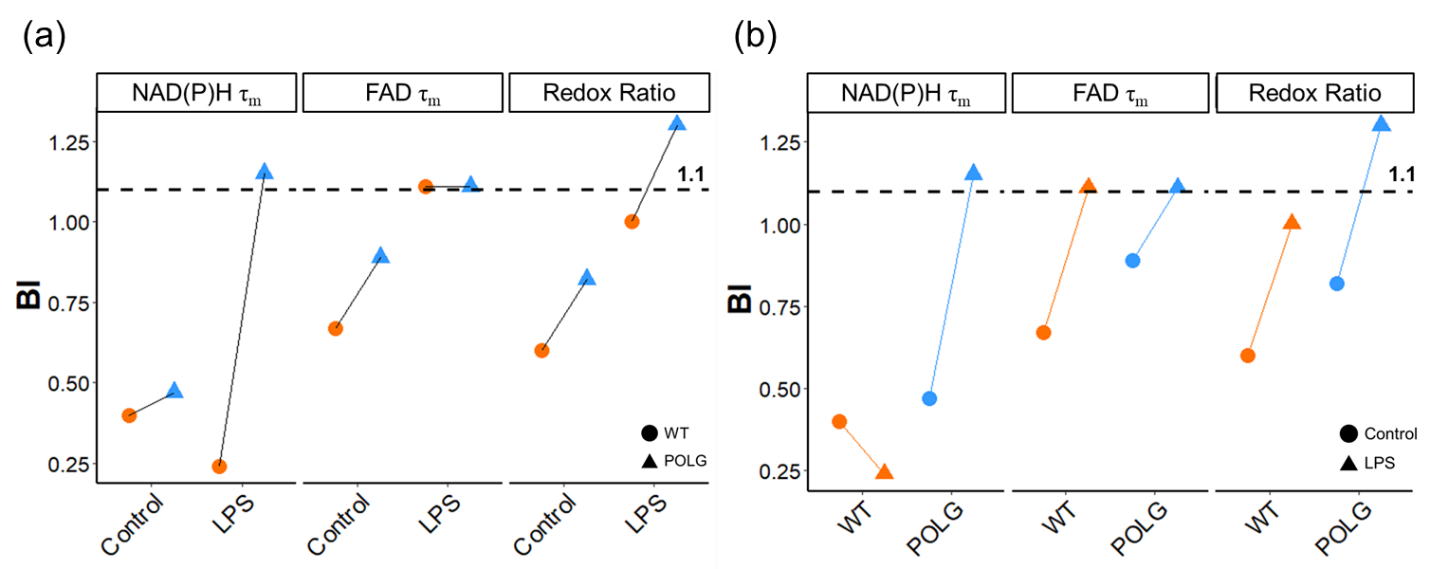

**Figure 2.** Metabolic heterogeneity analysis in control, and LPS-treated WT and POLG BMDMs for the combined dataset of three replicates. Comparison of BI of NAD(P)H *τ_m_*, FAD *τ_m_*, and optical redox ratio between (a) WT, and POLG BMDMs (b) Control and LPS-treatment.

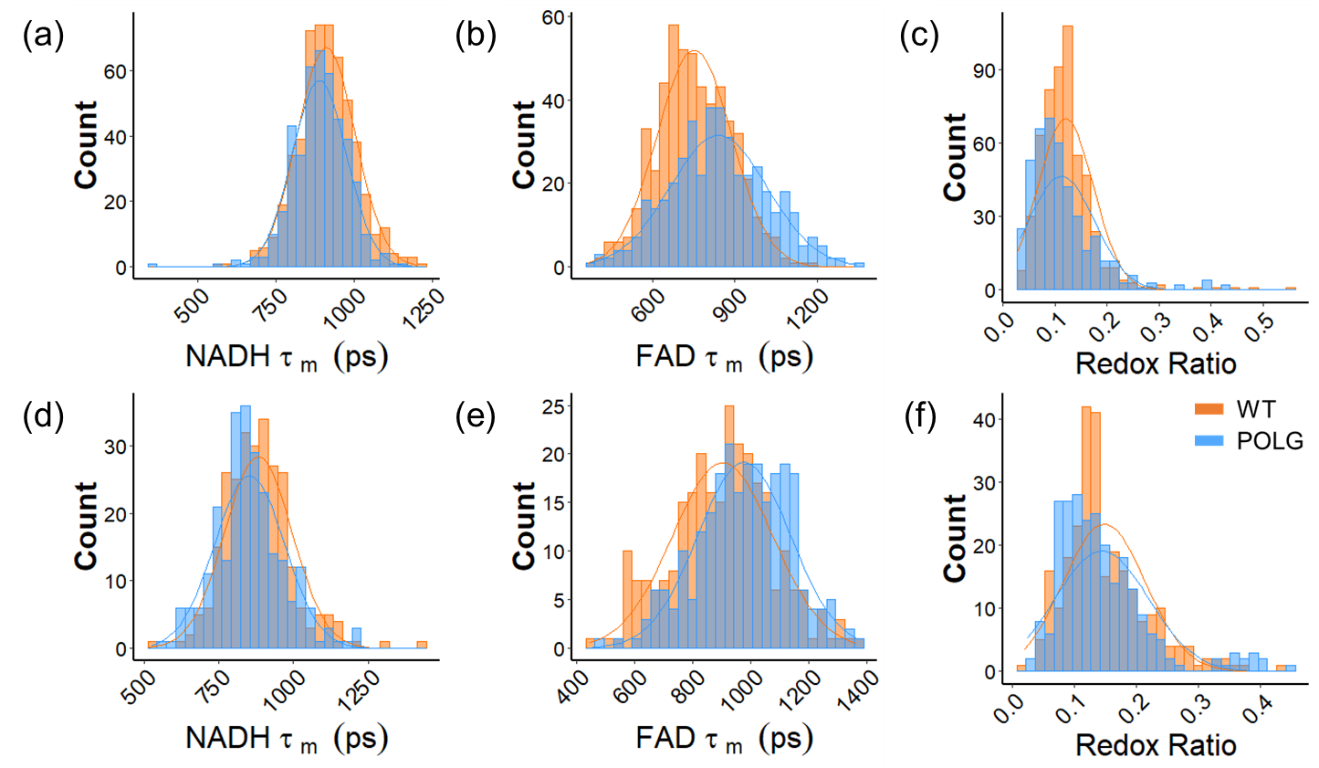

**Figure 3.** Histogram of NAD(P)H τ_m_, FAD τ_m,_ and redox ratio of WT and POLG BMDMs. Histogram of (a) NAD(P)H *τ_m_*, (b) FAD *τ_m_* (c) redox ratio of control WT and POLG BMDMs. Histogram of (d) NAD(P)H *τ_m_*, (e) FAD *τ_m_* (f) redox ratio of WT and POLG BMDMs with LPS treatment.

Table 6. Distribution analysis of NAD(P)H τ_m_, FAD τ_m_, and redox ratio for the cell population

|  | NAD(P)H *τ_m_* | | | FAD *τ_m_* | | | Redox Ratio | | |
| --- | --- | --- | --- | --- | --- | --- | --- | --- | --- |
|  | skewness | kurtosis | Std | skewness | kurtosis | Std | skewness | kurtosis | Std |
| Control WT | 0.065 | 0.603 | 92.3 | 0.111 | -0.167 | 136.1 | 2.818 | 15.715 | 0.053 |
| Control POLG | -0.689 | 3.540 | 86.1 | 0.069 | -0.271 | 178.1 | 1.940 | 5.552 | 0.064 |
| LPS WT | 0.720 | 2.265 | 113.0 | -0.002 | -0.202 | 173.1 | 1.091 | 1.560 | 0.066 |
| LPS POLG | 0.407 | 0.534 | 116.9 | -0.203 | -0.150 | 160.5 | 1.609 | 3.165 | 0.075 |

Std: standard deviation

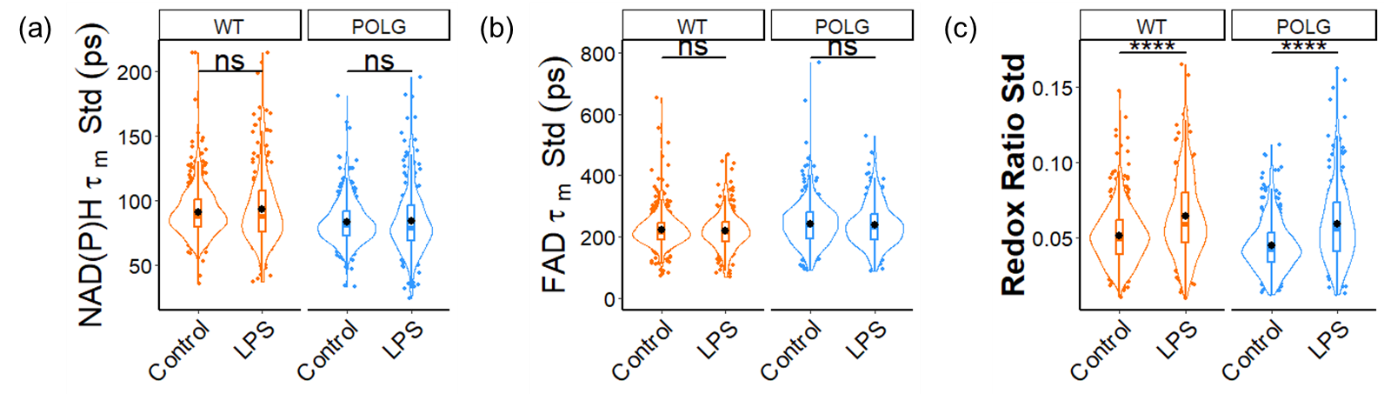

**Figure 4.** Intra-cellular standard deviation of (a) mean NAD(P)H fluorescence lifetime (b) mean FAD fluorescence lifetime and (c) optical redox ratio for untreated and LPS-treated WT and POLG BMDMs. *p < 0.05, and ****p < 0.0001 for two-sided student t-test. Statistics shown on plots for control-LPS comparisons for WT and POLG BMDMs.

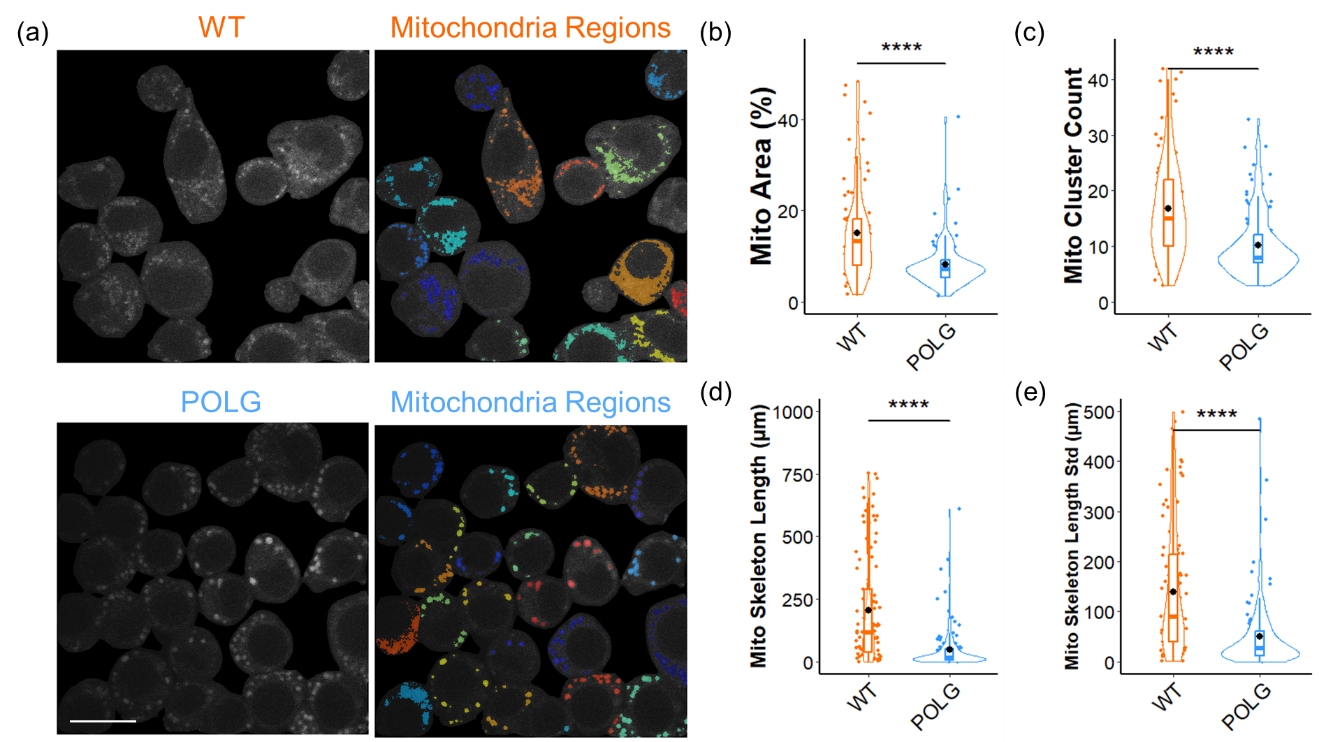

**Figure 5.** Mitochondria features of control WT and POLG BMDMs extracted from the NAD(P)H image captured with a 150x (1.35 NA) objective. (a) Representative NAD(P)H and segmented mitochondria images for WT and POLG BMDMs, scale bar = 15 *μm*. (b) Percentage of mitochondria area over whole cell area (c) Number of mitochondria clusters (d) Sum of the length of mitochondria skeleton (e) Standard deviation of length of mitochondria skeleton in WT and POLG macrophages. **** p < 0.0001 for two-sided student t-test.

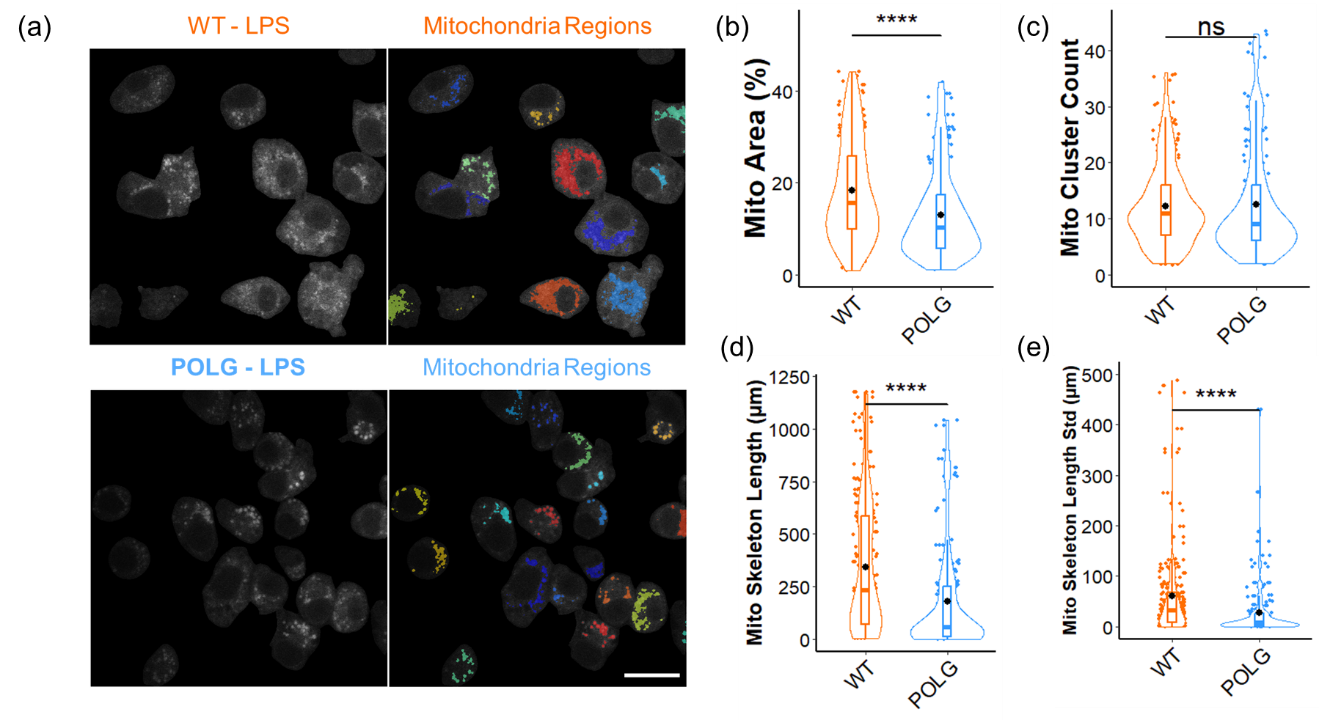

**Figure 6.** Mitochondria features of LPS-treated WT and POLG BMDMs extracted from the NAD(P)H autofluorescence image captured with a 100x (1.46 NA) objective. (a) Representative NAD(P)H and segmented mitochondria images for LPS-treated WT and POLG BMDMs, scale bar = 20 *μm*. (b) Percentage of mitochondria area over whole cell area (c) Number of mitochondria clusters (d) Sum of the length of mitochondria skeleton (e) Standard deviation of length of mitochondria skeleton in LPS-treated WT and POLG macrophages. **** p < 0.0001 for two-sided student t-test.

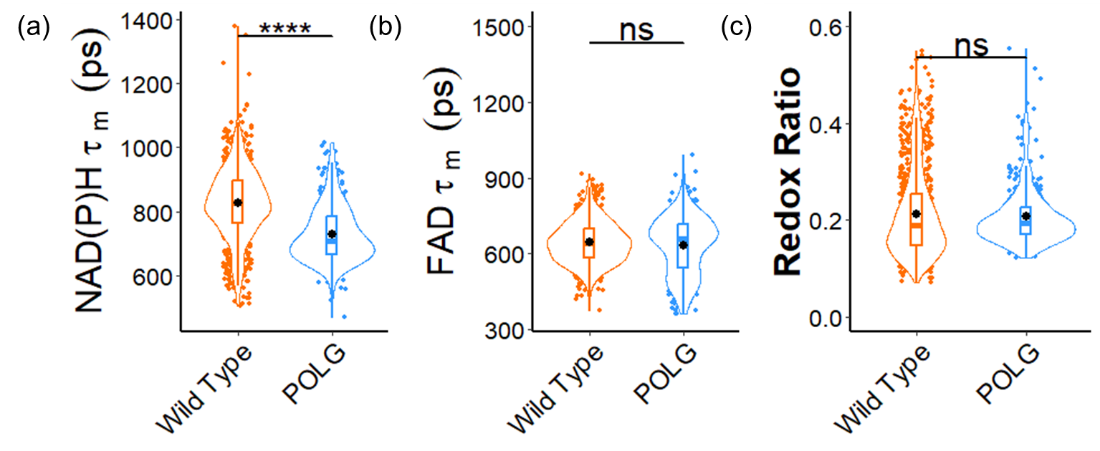

**Figure 7.** Autofluorescence lifetime imaging reveals different metabolic states between WT and POLG BMDMs of younger (7^th^ -10^th^) generations. (a) NAD(P)H *τ_m_* (b) FAD *τ_m_* (c) redox ratio of control WT and POLG BMDMs. ****p < 0.0001 for two-sided student t-test.
